# Transcriptomic analysis of *Picea abies* tissue culture reveals the impact of culture conditions and the presence of glucuronoxylan on extracellular lignin production

**DOI:** 10.1101/2024.06.11.598459

**Authors:** Ioanna Sapouna, Pramod Sivan, Francisco Vilaplana, Vaibhav Srivastava, Lauren Sara McKee

## Abstract

Tissue cultures are an important study model for woody plant tissue and can be used to study lignin biosynthesis. The greatest disadvantage of protocols based on extraction of lignin from wood biomass is the almost inevitable alteration of the native structure of lignin. Using a Norway spruce tissue culture with the ability to secrete monolignols into a liquid culture medium, fundamental aspects of lignin have been studied in the past, such as its structure, the enzyme activity related to its polymerization, and its interactions with a secondary cell wall hemicellulose. In this study, parameters that can induce monolignol production and secretion in the tissue culture are investigated via gene expression analysis. The impact of the composition of the solid growth medium, which was in some cases supplemented with xylan, was studied in depth through transcriptomic investigation. We find that the state (i.e. liquid or solid) and the xylan content of the medium can impact gene expression, although microscopic analysis suggests that cellular morphology is consistent. Extracellular lignin was collected from a formulation of liquid medium with the same composition as that used for cellular growth, which was previously presumed to be “non-inducing” of lignin biosynthesis. Chemical analysis of this lignin was performed using nuclear magnetic resonance spectroscopy and size exclusion chromatography, which revealed changes in its structure compared to the polymer produced in the previously developed “inducing” liquid medium. These experiments show that there is still much we do not understand about an oft-used tissue culture system, but show the way to a deeper understanding of the genetic control of lignin biosynthesis.

## Introduction

Norway spruce (*Picea abies* L. Karst) is one of the most economically significant trees in the world.^1^ Sweden is almost 70% covered in forest, of which approximately 40% comprises spruce.^2^ This makes the gymnosperm species a cornerstone of the Swedish forest ecosystem, as well as a foundational feedstock for the pulp and paper industry. The wood of gymnosperms is called softwood. Softwood composition has been mainly studied for the improvement of pulping processes and the final properties of wood-derived products. From a fundamental research point of view, softwood is an important representative of wood to study plant cell wall biosynthesis and architecture. For example, lignin, one of the three major components of wood, comprising approximately 27% of the dry softwood biomass^3^, consists almost solely of monolignols deriving from coniferyl alcohol (G-units).^4^ By comparison, in hardwood, lignin is composed of three major monolignols, making its structure more labile to extraction methods, and therefore more difficult to study.^4, 5^ The native structure of lignin is still keenly debated, mainly due to its variability depending on plant species, growing stage, and stress conditions.^4-8^ In addition, lignin extraction methods are known to alter the native structure, not only hindering our efforts to understand the architecture of the plant cell wall but also reducing the feasible options for utilising lignin.^3^ To improve our understanding of the unmodified structure of native lignin, cell cultures as models of wood tissue are an invaluable tool that enables insight into lignin biosynthesis and the interactions of lignin with polysaccharides. The monolignol components of the lignin polymer are thought to be produced via the phenylpropanoid biosynthetic pathway, which starts with deamination of phenylalanine and continues with subsequent modifications on the aromatic ring and aliphatic chain. This is regulated and controlled by enzymes, and leads to the synthesis of the three most common monolignols.^4, 9^ Recent studies have shown that there are other compounds with the ability to incorporate into the propagating lignin polymer that derive from outside of this pathway.^6^

A Norway spruce cell culture initiated by Simola et al. in 1985^10^ and maintained as callus culture on solid nutrient medium has been employed in previous studies to investigate lignin production^11^ and lignification-related enzyme activity, deriving from peroxidases and lacasses^12, 13^. According to the literature, 97% of the cells maintained on solid medium do not form a secondary cell wall, so no lignin deposition occurs.^14, 15^ However, the cells have the ability to produce extracellular lignin (ECL), when transferred into liquid medium suspension cultures that induce monolignol secretion.^16^ A recent paper theorised that extracellular vesicles (EVs) may potentially be involved in secreting packages of precursors and enzymes for lignin synthesis.^16^ In several studies ECL has been obtained and structurally characterised that confirmed that the product resembles the structure of so-called developmental lignin more than synthetic lignin.^11, 17-20^ There are also some differences between ECL and true cell wall lignin, and these have been attributed to the absence of a polysaccharide matrix, into which lignin is normally deposited during physiological lignification. To address this, in a previous study we added a secondary cell wall hemicellulose, xylan, into the nutrient medium of solid and suspension tissue cultures and investigated its role on monolignol production and the final structure of ECL.^21^ We concluded that cell growth and monolignol production are affected by the presence of xylan in the nutrient media, and that the structure of the ECL was affected as well, likely due to interactions with the xylan.

In this work we dig deeper into the factors that induce or affect monolignol secretion in the Norway spruce cell culture by, for the first time, analyzing lignin produced in suspension cultures in a theoretically “non-inducing” medium formulation. We used microscopy to visualize possible changes in the cells’ morphology, which may be caused by the different culture conditions and/or by the presence of xylan. Moreover, we performed an in-depth transcriptomic survey of gene expression in multiple different culture treatments, and preliminarily find that several pathways relating to lignin biosynthesis show differential expression in different culture treatments.

## Materials and Methods

### Materials

Chemicals for the preparation of nutrient media were purchased from Sigma-Aldrich, Germany and used without further purification. Absolute ethanol was purchased from VWR, Sweden. Purified beechwood xylan was purchased from Megazyme, Ireland. RNA was extracted with RNeasy plant mini kit purchased from Qiagen AB, Sweden. Quantification of extracted RNA was performed using a Qubit RNA HS assay kit, purchased from Thermo Fisher Scientific, Sweden.

### Culture media and treatments

The composition of nutrient media is described elsewhere.^13, 14^ As in previous work^21^, xylan was added into the solid and/or liquid nutrient medium in concentrations of 0.5 g/L and 1 g/L, before pH adjustment and autoclaving. In standard experiments, an “ECL non-inducing” medium is used for solid culture, and an “ECL-inducing” medium is used for suspension cultures. In some experiments, a liquid medium with the same composition as the “ECL non-inducing” one, used in the solid culture was also produced. The “ECL-inducing” liquid medium is marked with “i". Treatments were according to previous work and summarized in **Table 1**, together with sample nomenclature.

**Table 1:** Treatments as described in previous work.^21^ As an example of the sample nomenclature, 1→0.5i is a treatment in which cells pre-grown on 1 g/L xylan containing solid medium are transferred into 0.5 g/L xylan containing liquid medium. 0→0i is the reference.

|  | Xylan in liquid medium (g/L) |  |  |  |
| --- | --- | --- | --- | --- |
|  |  | 0 | 0.5 | 1 |
| Xylan in solid medium (g/L) | 0 | 0→0i \| 0→0 | 0→0.5i | 0→1i \| 0→1 |
|  | 0.5 | 0.5→0i | - | 0.5→1i |
|  | 1 | 1→0i \| 1→0 | 1→0.5i | 1→1i \| 1→1 |

### Maintaining the cell culture and inducing extracellular lignin production

As described in previous work, Norway spruce cells (Picea abies (L.) Karst., line A3/85) were maintained on solid medium and, when needed, transferred into liquid medium suspension cultures, in which monolignol secretion was induced.^14, 21^ Every three weeks, the cells were subcultured to fresh solid medium. The transfer into suspension cultures was performed two weeks after a subculture. The suspension cultures consisted of approximately 3 g cells (fresh weight) in 100 mL of liquid medium, in 500 mL flasks.

### Extracellular lignin collection and analysis

ECL was collected from the “non-inducing” liquid culture medium (0→0, 0→1, 1→0 and 1→1), by simple filtration of the suspension cultures with Miracloth (Merck Millipore), once it was visibly produced in the flasks, and while the cells were still of bright green colour. Several flasks were used per treatment to ensure a sufficient ECL amount for chemical analyses.

### Size exclusion chromatography

Molecular weight characterization experiments of ECL were performed using a gel permeation chromatography (GPC) system from Waters (Waters Sverige AB, Sollentuna, Sweden) consisting of a Waters-515 high pressure liquid chromatography (HPLC) pump, a 2707 autosampler, a 2998 photodiode array detector (UV detector) operated at 254 nm and 280 nm, and a 2414 refractive index (RI) detector. Waters Ultrastyragel HR4, HR2 and HR0.5 (4.6 × 300 mm) solvent efficient columns were used, connected in series with a Styragel guard column, set to 35 °C. Calibration was performed using polystyrene standards with molecular weights from 176 kDa to 370 Da, using the data from the 254 nm channel of the UV detector. Flow rate of tetrahydrofuran (THF) eluent was 0.3 mL/min. The ECL samples were acetylated before the analysis. The protocol is described in previous work.^3^ In short, 2 mg of sample was mixed with 200 μL pyridine:acetic anhydride solution (1:1 vol.), at room temperature, overnight. Pyridine was removed by addition of cold toluene:methanol solution (1:1 vol.), and evaporating under nitrogen flow. The addition of toluene:methanol solution was repeated at least four times.

### Nuclear magnetic resonance spectroscopy

A Bruker 400 MHz DMX instrument was used (Bruker Corporation, Billerica, MA, USA), with a multinuclear inverse Z-grad probe. The pulse sequence used for the HSQC experiments was hsqcetgpsi. The pulse length was optimized at 9.2 s with D1 of 1.49 s. 176 scans were used per sample.

### Cell morphology analysis by optical microscopy

For lignin localization, cells mounted on a glass slide were subjected to Wiesner staining (one volume of 37 N HCl, two volumes of 3% phloroglucinol in ethanol), covered with cover slip^22^, and observed under microscope. For fluorescence microscopy, lignin autofluorescence was measured at green excitation (470 nm) and emission at 525nm^23^ while for the cell viability test, cells were stained with 10 μg/mL propidium iodide followed by observation at green excitation (405-490 nm) and emission at 610-660nm^24^. Both bright field and fluorescent images were taken using a Leica Dmi8 microscope (Leica Microsystems, Germany) with a color DCF7000 T camera.

### Transcriptomic analysis

### Collection of cell samples and extraction of RNA

Cell samples were collected from pooled, filtered cells in Eppendorf tubes. The cell samples were immediately frozen with liquid nitrogen and kept at -80 °C until RNA extraction. The ECL-containing filtrate was centrifuged at 17,000 g for 40 min, using a JA10 rotor in an Avanti J-265 XP centrifuge (Beckman Coulter). The pelleted ECL was washed twice with Milli-Q water and freeze-dried before chemical analysis.

Using an RNeasy plant mini kit, RNA was extracted from cells collected and stored as described above. The kit was used according to the manufacturer’s instructions. Briefly, the frozen cells were ground to a fine powder using mortar and pestle. Approximately 100 mg of the cells were used lysed with the RLT buffer. The lysate was transferred to a QIAshredder spin column, and centrifuged at room temperature. Absolute ethanol was added to the RNA-containing pass-through column and the mixture was transferred to an RNeasy spin column. The binding membrane of the column was washed first with the RW1 buffer and then with the RPE buffer. Finally, the RNA bound on the membrane of the column was collected with RNAse-free water, with two washings. The RNA samples were frozen in liquid nitrogen and kept at -80 °C until sequencing.

Cells were collected from reference solid culture and 1, 2, and 3 weeks after they were subcultured to xylan-containing solid medium. In addition, cells were collected as described in the previous section, five days after they were transferred into suspension cultures and after ECL was collected. All cell samples collected are summarized in **Table 2**.

**Table 2:**
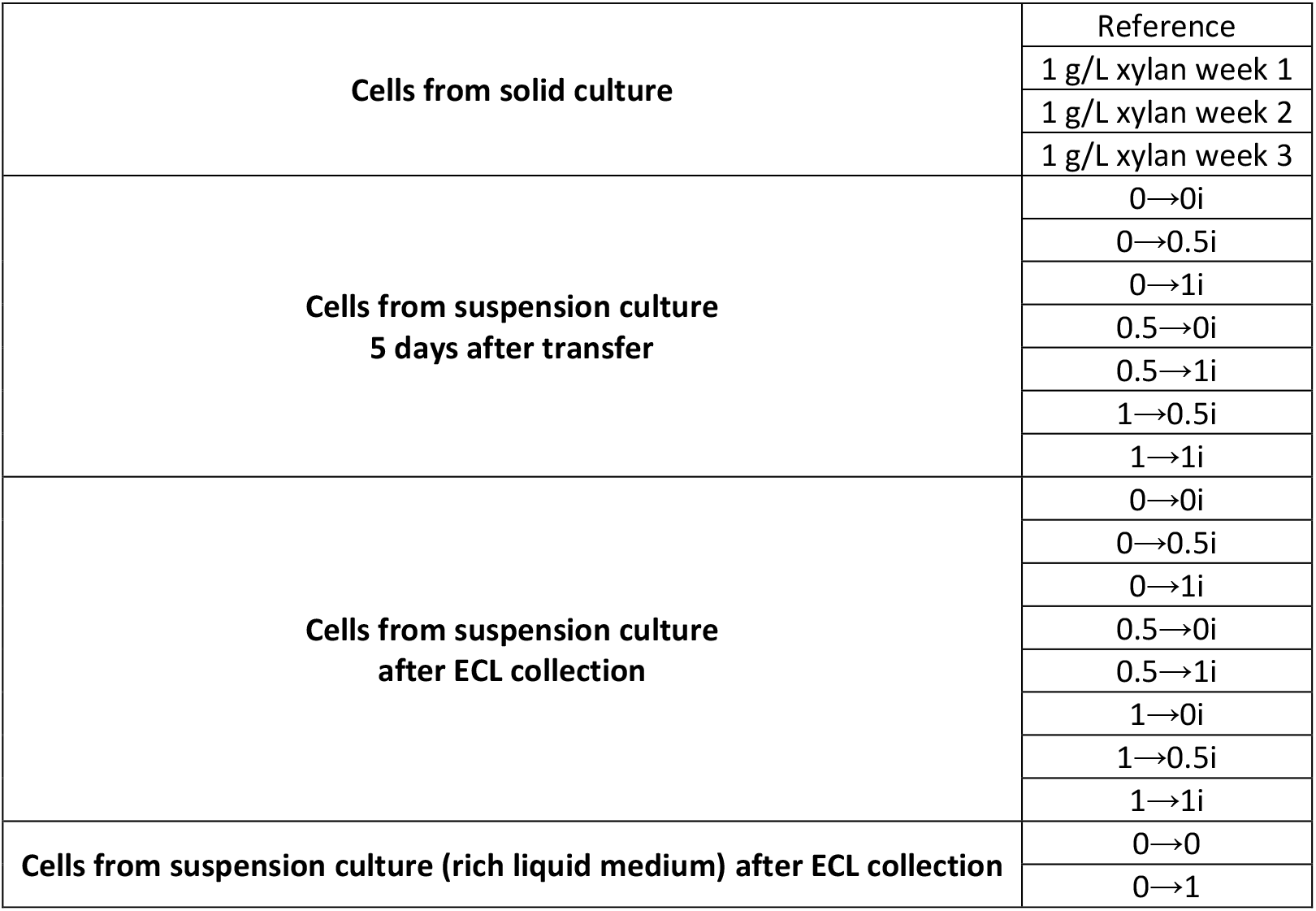
Summary of the cell samples collected for RNA extraction. Nomenclature of the samples is described in.

### RNA sequencing

RNA sequencing was performed by the National Genomics Infrastructure at SciLife lab, Stockholm, Sweden. The protocol for library preparation with the Illumina TruSeq Stranded mRNA kit (Illumina Cat no: 20020595, Illumina, USA) was adapted for automation on the Agilent NGS Bravo workstation (Agilent Technologies, USA) for a 96-well plate format. All reagents used for these steps are supplied as components of the Illumina TruSeq Stranded mRNA kit unless otherwise stated. Library preparation with the Illumina TruSeq Stranded mRNA kit (Illumina Cat no: 20020595, Illumina, USA) comprised the following: quality checks on total RNA; selective purification of mRNA from total RNA; synthesis of cDNA; adapter ligation; amplification; quantitation of libraries for concentration estimates. The concentration of RNA was estimated using a Qubit 3.0 (Cat no: Q33238, RNA HS Assay Kit; Cat no:Q32855, ThermoFisher Scientific), and the quality of RNA based on the RNA Integrity Number was estimated using either the 2100 BioAnalyzer (Cat no: G2939BA; RNA 6000 Nano Kit; Cat no: 5067-1511. Agilent), 5200 Fragment Analyzer (Cat no: M5310AA; High Sensitivity RNA (15nt) DNF-472 kit; Cat no: DNF-472-0500. Agilent), or Caliper GX Touch instrument (Cat no: CLS137031, RNA Assay reagent Kit; Cat no: CLS960010. PerkinElmer). The subsequent steps of library preparation were all carried out on the Agilent NGS Bravo workstation in 96-well plates following the instructions for the Illumina TruSeq Stranded mRNA kit (Illumina Cat no: 20020595, Illumina, USA). mRNA was purified from 300-800 ng of total RNA through selective-binding on poly dT-coated beads, and fragmented using divalent cations under elevated temperature. cDNA was synthesized from the resulting fragments by adding reverse transcriptase SuperScript II Reverse Transcriptase (Cat no: 1808004. ThermoFisher Scientific). This step was followed by bead clean-up with the AMPure XP solution (Cat no: A1905B. Beckman Coulter) to selectively retain fragments of desired lengths. cDNA was then subjected to 3’ adenylation, which was followed by adapter ligation to the 3’ adenylated end of the fragment. The fragments with ligated adapters were once again cleaned-up on AMPure XP beads to remove non-ligated adapters, followed by PCR to amplify these fragments. The PCR products were purified by binding to AMPure XP beads, washed with 80% ethanol and eluted in EB (Cat no: 19086. Qiagen). The quality of the adapter-ligated libraries was checked on the BioAnalyzer High Sensitivity DNA kit (Cat no: 5067-4626. Agilent) or Caliper GX LabChip GX/HT DNA high sensitivity kit (Cat no: CLS760672, PerkinElmer) and concentration by Quant-iT DNA High sensitivity kit (Cat no: Q33232. ThermoFisher).

### Transcriptomic data analysis

Analysis of transcriptomic data was performed by the National Bioinformatics Infrastructure of Sweden, as follows. Data QC was performed with FastQC v0.11.9.^25^ Raw reads were trimmed for low quality sequences, and adaptor sequences were removed using the Fastp program.^26^ The resulting reads were fed to Salmon v1.4.0^27^ to quantify transcripts against the reference spruce transcriptome downloaded from https://plantgenie.org database. Transcript-level counts from Salmon were summarized into a matrix by tximport R package v1.20.^28^ For the differential expression analysis, gene expression counts were normalized with the R package DESeq2 v1.32.2.^29^ For PCA analysis used in data consistency checks, gene counts were transformed into log2 scale by applying regularized logarithm approach using the rlog function provided in DESeq2 package. Pathway analysis of differentially expressed genes was performed using the KEGG (Kyoto Encyclopedia of Genes and Genomes) Automatic Annotation Server (https://www.genome.jp/kegg/kaas/).

## Results and Discussion

### Changes in cultivation condition lead to the production of ECL

The Norway spruce tissue culture cells are maintained on solid medium, where they do not form a secondary cell wall, as evidenced by optical microscopy experiments (**Figure 1**). In addition, the primary walls of the cells are not lignified, observed by the absence of staining with the Wiesner test (**Figure 1**, top row). The reagent used in the Wiesner test is known to bind to aldehyde groups in lignin, which can be present both as end groups and within the lignin polymer.^30^ Since lignin does not polymerize inside the plasma membrane^31^, the staining observed in cells collected from solid medium with xylan provision (**Figure 1**, 0.5 g/L xylan and 1 g/L xylan) was attributed to the presence of monolignols. In the xylan treatments, there was no increase in the number of differentiated cells observed, suggesting that xylan does not induce secondary cell wall formation in this culture, but it may have a role in inducing monolignol production. Some microscopy images suggest that monolignols may be secreted from cells while growing on xylan-supplemented plates (**Figure 1**, 1 g/L xylan) but we find no evidence of monolignol accumulation or lignin formation in the agar (**Figure S1**).

**Figure 1:**
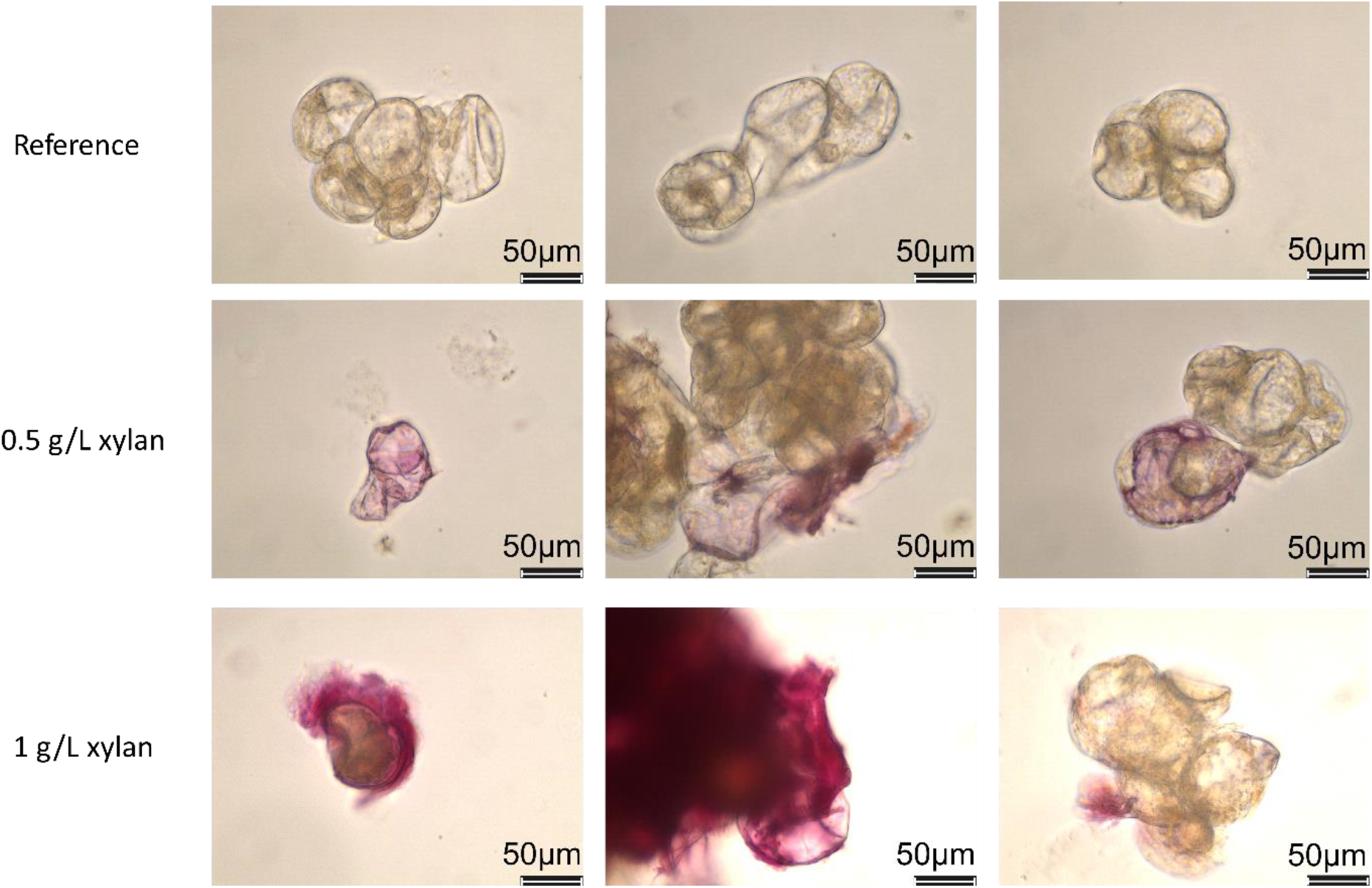
Wiesner test as captured by optical microscopy. The cells were collected from solid cultures with the described xylan provision. Three images of each treatment are shown, to represent the diversity in the observed cell population. Magnification is 40x and the scalebars are 50 μm.

After two weeks of pre-growth in solid culture, the cells were transferred into suspension culture, to induce monolignol secretion, which was until now thought to be a result of changes in the composition of the medium.^10^ Light and temperature conditions, which are also known to promote lignin formation, were the same for all solid and suspension cultures. To confirm whether the medium composition is indeed the inducing factor in monolignol secretion and ECL production, we created a liquid medium with the same composition as that used for the solid culture, a medium referred to as “non-inducing” from now on. Cells were inoculated from solid culture into the “non-inducing” liquid, creating the 0→0, 0→1, 1→0, and 1→1 treatment conditions. Interestingly, the cells produced ECL in all cultivation flasks, at approximately the same time as it occurred for the “ECL-inducing” liquid medium used in previous work (13-20 days).^21^

Composition analysis of the ECL produced in the suspension cultures with the “non-inducing” medium showed that the polymer was larger in size than that produced in the “ECL-inducing” medium used in previous work.^21^ The number average molecular weight (Mn) varied between 2000-2800 Da, compared to 1700-1900 Da for the ECL collected from the “ECL-inducing” liquid medium (**Table S1**). In addition, ECL in this work had a higher β-O-4’ content, ranging between 30-40% compared to 26-31% in our previous study^21^. The NMR analysis was conducted on two ECL samples collected separately, and the integral values were similar, showing good reproducibility of the effect of the treatment on the ECL production (**Table S2**). Interestingly, the ECL product in this study had a very different appearance compared to the one collected in the previous work. More specifically, the ECL product collected here had a hydrogel-like texture. After freeze-drying, it was not possible to dissolve it in DMSO-*d6*. An extraction with 80% ethanol was performed overnight, at room temperature, to obtain a more soluble, lignin-enriched fraction for analysis. The ECL-containing product most probably consists of a large amount of pectins and xyloglucan, which could cause the hydrogel-like texture. In previous work, the ECL product from the “ECL-inducing” liquid medium contained these polysaccharides^19^, and we think that they could be secreted in the “non-inducing” system as well. We speculate that the polysaccharides in this new polymerization environment might differ in composition and amount, creating a different matrix for the monolignol polymerization, resulting in the observed differences between the ECL collected from the “ECL-inducing” and “non-inducing” liquid media.

In our earlier work, we observed an increased ECL production in cultures with 0.5 or 1 g/L xylan supplemented into the solid culture medium. Here, we have investigated the impact of xylan on cells grown in xylan-containing solid culture then transferred into the “non-inducing” liquid medium containing xylan. Although further work is required to quantify the amount of ECL produced, optical microscopy images of cells collected from suspension cultures with the “ECL-inducing” liquid medium confirmed monolignol production inside the cells, irrespective of xylan provision (**Figure S2**). The same was observed for the suspension cultures with the “non-inducing” medium: in **Figure 2**, autofluorescence of monolignols produced inside the plasma membrane in cells from “non-inducing” liquid medium is presented.

**Figure 2:**
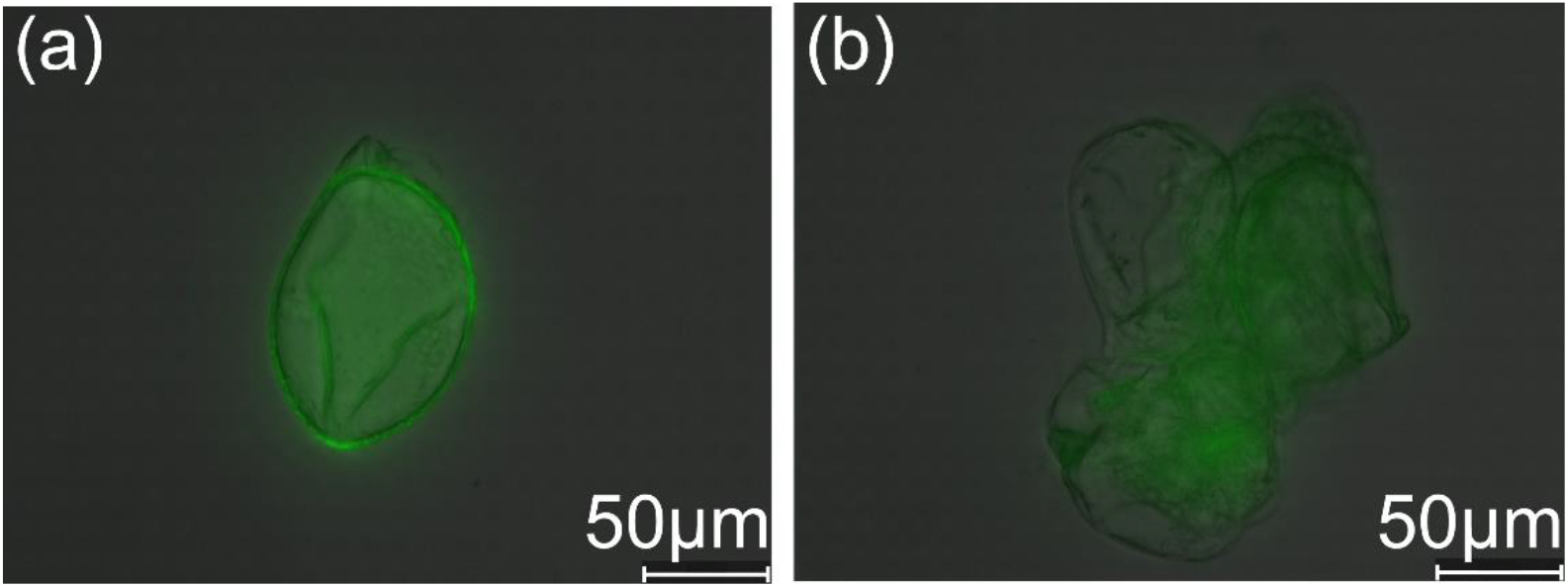
Overlay of autofluorescence and bright background of cells collected from “non-inducing” liquid medium after ECL was collected. The treatments are (a) 0→0 and (b) 1→1. Magnification of the images is 40 x and the scale bar is 50 μm.

### Differential gene expression in treatments with xylan provision

In typical experiments^21^, cells were pre-cultivated on solid medium and then transferred into an “ECL-inducing” liquid medium into which monolignols were secreted, and spontaneously assembled into the ECL polymer. We extracted total RNA from cells in the suspension culture, collected on the same day that ECL was collected, and submitted this for transcriptomic analysis. Transcript abundance was quantified for 44838 genes. Samples were compared in a pair-wise manner to determine the fold-change in expression between culture conditions. A difference in transcript abundance between two samples indicates up- or down-regulation of the expression of a certain gene, and this can be expressed in relative terms that compare two samples. A 2.5-fold difference in expression level is often considered to be significant. In our dataset, a large number of genes displayed a 2.5-fold difference in expression between most pairs of samples that were compared, and so genes were also identified that showed a more significant 5- or 7.5-fold change in expression between samples, to further narrow down the list of potentially important genes. The number of genes differentially expressed in relevant pairs of culture conditions gives some insight into how strongly affected the cells are by the culture condition. For example, the effect of the addition of xylan into this liquid medium can be seen in the comparisons in **Table 3**. The largest change (number of genes showing differential expression of at least 5- or 7.5-fold) is observed for the comparison between 1→1i and 1→0i. This suggests that gene expression is strongly affected by the presence vs absence of xylan in the “ECL-inducing” liquid medium when cells are pre-grown on xylan-containing solid medium.

**Table 3:** Differential gene expression in treatments with “ECL-inducing” liquid medium, with or without xylan provision, as indicated. ‘Number of genes’ in a comparison of sample A vs sample B is the number of genes showing a log2fold change in expression of at least 2.5x, 5x or 7.5x, as indicated. An X-fold difference in expression counted here can be an upregulation of either sample in the comparative pair.

| Sample A | Sample B | Number of genes |  |  |
| --- | --- | --- | --- | --- |
|  |  | 2.5-fold | 5-fold | 7.5-fold |
| 0→1i | 0→0i | 43697 | 71 | 5 |
| 1→1i | 1→0i | 40424 | 594 | 65 |
| 1→0.5i | 1→0i | 42866 | 232 | 25 |

The effect of the addition of xylan into the solid medium can be seen in **Table 4**. With the exception of 0.5→1i compared to 0→1i, which shows the largest difference, there is a trend observed with greater differences found when comparing samples with vs without xylan in the solid medium.

**Table 4:** Differential gene expression in treatments with “ECL-inducing” liquid medium, with or without xylan provision, as indicated.

| Sample A | Sample B | Number of genes |  |  |
| --- | --- | --- | --- | --- |
|  |  | 2.5-fold | 5-fold | 7.5-fold |
| 0→0.5i | 1→0.5i | 44193 | 133 | 13 |
| 0.5→0i | 0→0i | 43860 | 21 | 0 |
| 0.5→1i | 0→1i | 39649 | 566 | 35 |
| 1→0i | 0→0i | 43692 | 39 | 2 |
| 0→1i | 1→1i | 40579 | 364 | 19 |
| 0→0.5i | 1→0.5i | 44193 | 133 | 13 |

### Differential gene expression in ECL “non-inducing” liquid medium

Although the impact of xylan in the solid and liquid media is illuminating, the largest differences in gene expression were in fact observed when gene expression in the “non-inducing” liquid was compared to that in the “ECL-inducing” liquid medium. Relevant comparisons are presented in **Table 5**. The first comparison in the table clearly shows the impact of agitation on gene expression. The same chemical composition was used for the solid and liquid media and, because the light conditions were also the same, the only difference between 0→0 cells collected from the “non-inducing” liquid and reference cells collected from the xylan-free (reference) solid medium was the agitation introduced in the suspension culture, which is necessary for oxygenation of the liquid medium. Although the cells do not produce or secrete monolignols in the reference solid medium (**Figure 1**, and **Figure S1**), they do so when introduced into suspension cultures with the same medium composition, as discussed in the previous section, and it is striking to see that over 1500 genes show differential expression in these two conditions (5-fold level of difference). This pronounced impact of the different liquid medium composition is obvious in the comparisons 0→0 to 0→0i and 0→1 to 0→1i.

**Table 5:** Changes in gene expression in treatments with “non-inducing” and “ECL-inducing” liquid medium, with or without xylan provision, as indicated.

| Sample A | Sample B | Number of genes |  |  |
| --- | --- | --- | --- | --- |
|  |  | 2.5-fold | 5-fold | 7.5-fold |
| 0→0 | Reference solid medium | 37539 | 1674 | 350 |
| 0→0 | 0→0i | 37015 | 1858 | 407 |
| 0→1 | 0→1i | 36806 | 1927 | 397 |
| 0→1 | 0→0 | 42624 | 4584 | 9 |

### Changes in expression level for genes in specific functional groupings

A KEGG pathway analysis was performed to determine the metabolic processes that may be impacted by the differential gene expression we have seen in these experiments. We have performed a detailed analysis of pathway involvement for genes showing differential expression between samples 1→1i and 1→0i, so that we may understand the changes that occur when xylan is added into the “ECL-inducing” liquid medium. In comparing these samples, a number of genes in certain key pathways show differential gene expression levels. In **Table 6** we highlight observed gene expression differences in pathways relating to lignin biosynthesis (marked with an asterisk on the table). We also highlight the other pathways that contain the highest number of genes showing upregulation in 1→1i compared to 1→0i, as this may illuminate other metabolic changes induced by the addition of 1 g/L xylan into the liquid medium, where monolignols are secreted. It is clear that, in addition to the changes in lignin biosynthesis (KEGG groupings 00940, 00400, 000360), there are changes that seem to affect general metabolism (04075, 00941, 04016, 00592), carbohydrate metabolism (00500, 00010), and stress responses (04626, 05208). Stress response has previously been correlated with ECL production (ref).^4-8, 16^ Indeed, we see some evidence of increased expression of stress-related genes such as chitinases in plate-growth conditions where xylan is provided, perhaps suggesting that the hemicellulose causes some form of stress response. However, although there are changes in the relative expression levels of various stress-related genes in our experimental systems, the data do not show a clear enough trend that one particular variable can be presumed to be the stressor underlying changes in lignin production.

**Table 6:** Pathways in which genes are differentially expressed in two culture conditions. Pathways relating to lignin biosynthesis are marked with an asterisk (*).

| Pathway KEGG identifier | Pathway description | Number of genes upregulated in 1→1i compared to 1→0i | Number of genes downregulated in 1→1i compared to 1→0i |
| --- | --- | --- | --- |
| 04075 | Plant hormone signal transduction | 10 | 10 |
| 00940* | Phenylpropanoid biosynthesis* | 9 | 6 |
| 00941 | Flavonoid biosynthesis | 8 | 4 |
| 04016 | Mitogen-activated protein kinase (MAPK) signaling pathway (cellular response to stimuli) | 8 | 5 |
| 04626 | Plant-pathogen interaction | 8 | 7 |
| 00500 | Starch and sucrose metabolism | 8 | 5 |
| 00010 | Glycolysis/Gluconeogenesis | 7 | 3 |
| 00620 | Pyruvate metabolism | 7 | 2 |
| 00400* | Phenylalanine, tyrosine and tryptophan biosynthesis* | 6 | 0 |
| 00592 | alpha-Linolenic acid metabolism | 6 | 0 |
| 05208 | Relating to reactive oxygen species | 6 | 4 |
| 00360* | Phenylalanine metabolism* | 4 | 0 |
| 00520 | Amino sugar and nucleotide sugar metabolism | 4 | 5 |

Furthermore, we have performed pathway analysis for genes showing differential expression between samples 0→0 vs 0→0i, and between samples 0→1 vs 0→1i. In these pairs, the concentration of xylan in the liquid medium is the same in both samples being compared, but the medium composition differs. This comparison will allow us to isolate the differences in gene expression arising from the use of the “ECL-inducing” vs the “non-inducing” liquid medium formulations. We observed that the ECL-containing product from the supposedly “non-inducing” liquid medium had a different appearance, likely due to a high concentration of carbohydrates, and that the ECL had different chemical properties than that collected from the “ECL-inducing” liquid medium. In **Table 7** we show the KEGG categories with the greatest number of genes with differential expression in the xylan-free samples (0→0 vs 0→0i), and in **Table 8** we present the same for the xylan-supplemented samples (0→1 vs 0→1i). In both, we also show the number of genes with differential expression in pathways relevant to lignin biosynthesis. As above in **Table 6**, the pathways highlighted in these comparisons relate to general metabolism, lignin biosynthesis, and stress response. No clear trends are however apparent for the differential (or inducible) expression of specific genes.

**Table 7:**
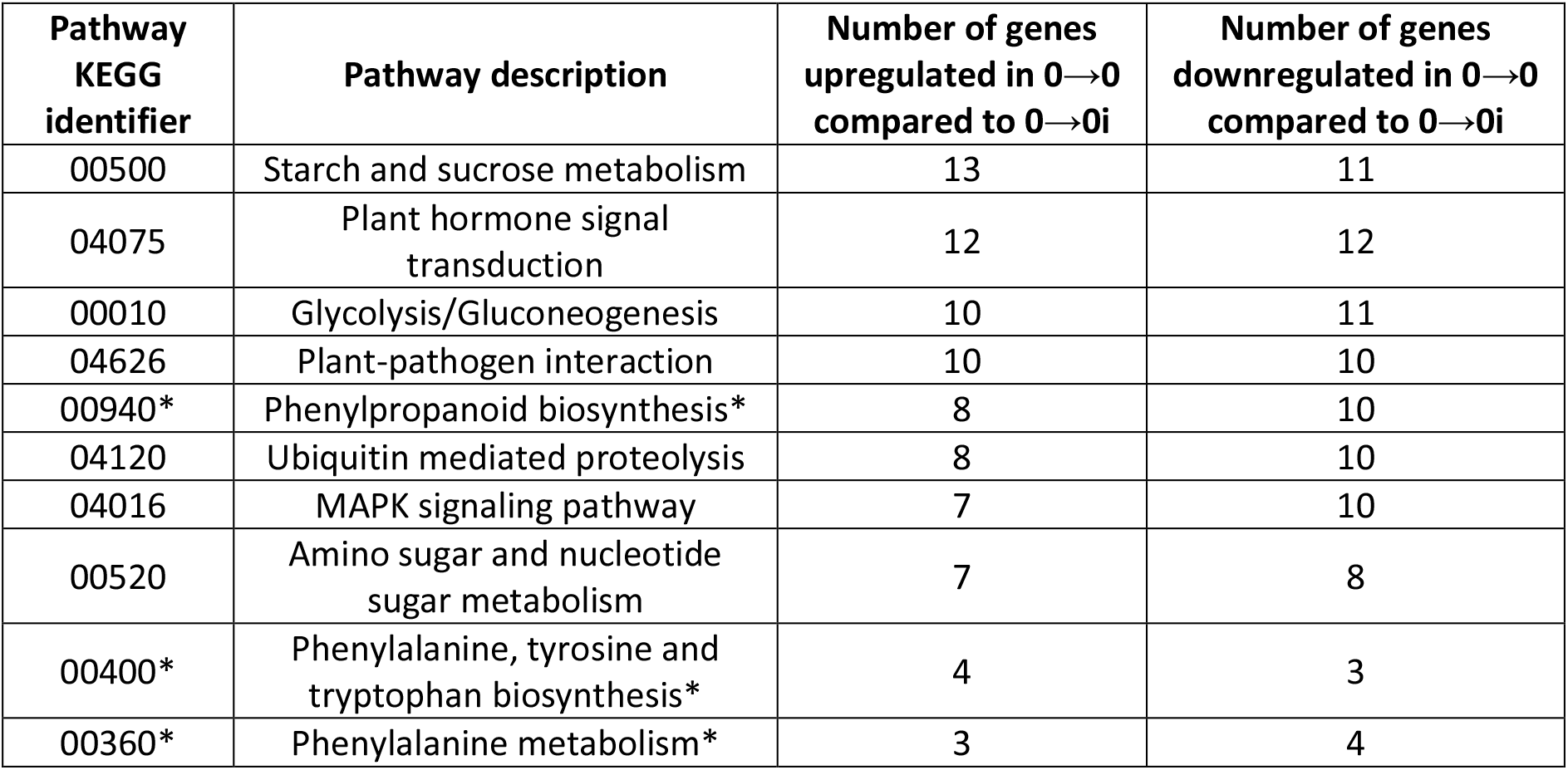
Pathways in which genes are differentially expressed in two culture conditions. Pathways relating to lignin biosynthesis are marked with an asterisk (*).

| Pathway KEGG identifier | Pathway description | Number of genes upregulated in 0→0 compared to 0→0i | Number of genes downregulated in 0→0 compared to 0→0i |
| --- | --- | --- | --- |
| 00500 | Starch and sucrose metabolism | 13 | 11 |
| 04075 | Plant hormone signal transduction | 12 | 12 |
| 00010 | Glycolysis/Gluconeogenesis | 10 | 11 |
| 04626 | Plant-pathogen interaction | 10 | 10 |
| 00940* | Phenylpropanoid biosynthesis* | 8 | 10 |
| 04120 | Ubiquitin mediated proteolysis | 8 | 10 |
| 04016 | MAPK signaling pathway | 7 | 10 |
| 00520 | Amino sugar and nucleotide sugar metabolism | 7 | 8 |
| 00400* | Phenylalanine, tyrosine and tryptophan biosynthesis* | 4 | 3 |
| 00360* | Phenylalanine metabolism* | 3 | 4 |

**Table 8:** Pathways in which genes are differentially expressed in two culture conditions. Pathways relating to lignin biosynthesis are marked with an asterisk (*).

| Pathway KEGG identifier | Pathway description | Number of genes upregulated in 0→1 compared to 0→1i | Number of genes downregulated in 0→1 compared to 0→1i |
| --- | --- | --- | --- |
| 04120 | Ubiquitin mediated proteolysis | 14 | 6 |
| 04075 | Plant hormone signal transduction | 12 | 16 |
| 00500 | Starch and sucrose metabolism | 12 | 12 |
| 00010 | Glycolysis/Gluconeogenesis | 11 | 12 |
| 00710 | Carbon fixation in photosynthetic organisms | 10 | 4 |
| 04016 | MAPK signaling pathway | 10 | 9 |
| 00520 | Amino sugar and nucleotide sugar metabolism | 8 | 6 |
| 04626 | Plant-pathogen interaction | 8 | 11 |
| 00270 | Cysteine and methionine metabolism | 8 | 12 |
| 00330 | Arginine and proline metabolism | 8 | 4 |
| 04141 | Protein processing in endoplasmic reticulum | 8 | 6 |
| 00940* | Phenylpropanoid biosynthesis* | 6 | 11 |
| 00400* | Phenylalanine, tyrosine and tryptophan biosynthesis* | 6 | 2 |
| 00360* | Phenylalanine metabolism* | 3 | 2 |

## Conclusions

In this study, it was possible to examine in depth the parameters that affect monolignol production in different tissue culturing conditions. Spruce cells that are maintained on solid culture medium do not produce monolignols, but when they are transferred into suspension culture with liquid medium of the same “non-inducing” composition, extracellular lignin (ECL) is formed. The structure of this ECL differed compared to the ECL produced in previous work, in the so-called “ECL-inducing” liquid medium. The impact of xylan on monolignol production was also investigated and it was shown with optical microscopy experiments that xylan provision in the solid medium can apparently trigger monolignol production, but likely not secretion or polymerization. The effect of both composition of the medium (liquid vs solid) and the provision of xylan on the lignin biosynthetic pathway was studied with transcriptomic analysis. The analysis indicates that several genes relating to lignin biosynthesis, and the biosynthesis of lignin precursors, are differentially expressed in different culture conditions, as well as several stress-related pathways and general metabolic pathways. The extensive dataset we have generated is a treasure-trove of important information as to how lignin production in cell suspension cultures might be impacted by cultivation conditions and the provision of exogenous carbohydrates. Researchers in the fields of lignin biosynthesis and/or plant cell suspension cultures are encouraged to use the dataset to seek additional perspectives on their own work.

## Supporting information

Data supporting the main manuscript

Raw RNASeq data

## Acknowledgements

The authors would like to acknowledge funding from the Knut and Alice Wallenberg Foundation through the Wallenberg Wood Science Centre. The authors are grateful to the Anna och Nils Håkansson and C.F. Lundström foundations for funding part of the RNA sequencing and data analysis experiments. In addition, the authors acknowledge support from the National Genomics Infrastructure in Stockholm funded by Science for Life Laboratory, the Knut and Alice Wallenberg Foundation, and the Swedish Research Council, and SNIC/Uppsala Multidisciplinary Center for Advanced Computational Science for assistance with massively parallel sequencing and access to the UPPMAX computational infrastructure. Lastly, the authors would like to acknowledge support from the National Bioinformatics Infrastructure Sweden. Specifically, we are grateful for help and advice received from Elísabet Einarsdóttir at NGI and Adnan Niazi from NBS.

## Data availability statement

The full dataset of gene transcript abundance is provided as a supplementary spreadsheet. Please cite this pre-print if you use the data in your own publications.

