## Supplementary material for "Transcriptomic analysis of *Picea abies* tissue culture reveals the impact of culture conditions and the presence of glucuronoxylan on extracellular lignin production": Data supporting the main manuscript

### Supporting Information

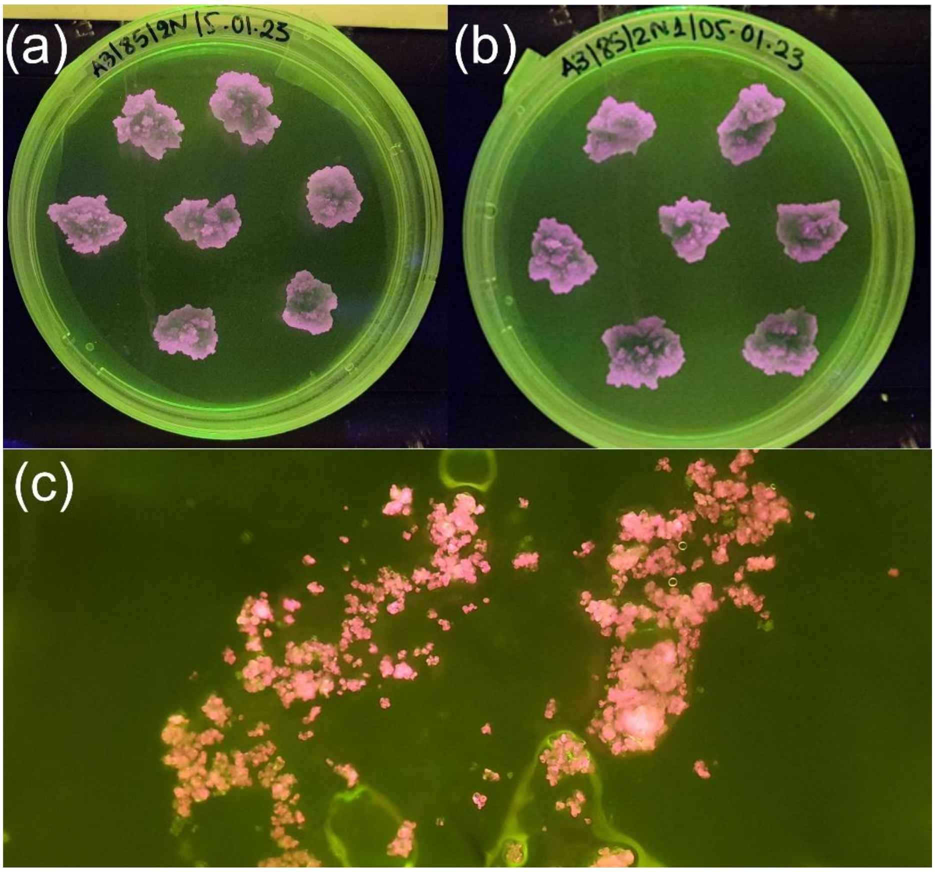


**Figure S1**: Plate with (a) reference solid culture, (b) solid culture with 1 g/L xylan and (c) zoom in on cells from (b) after the callus was removed. The picture was taken under green fluorescent light, which would show lignin/monolignols coloured pink. In (c) it is clearly shown that there is no coloration on the solid medium, but only in the cells, indicating no monolignols are released into the solid medium. The stain seemingly released from some cells in **Figure 1** is probably caused by breaking of the membranes during handling and chemical treatment.

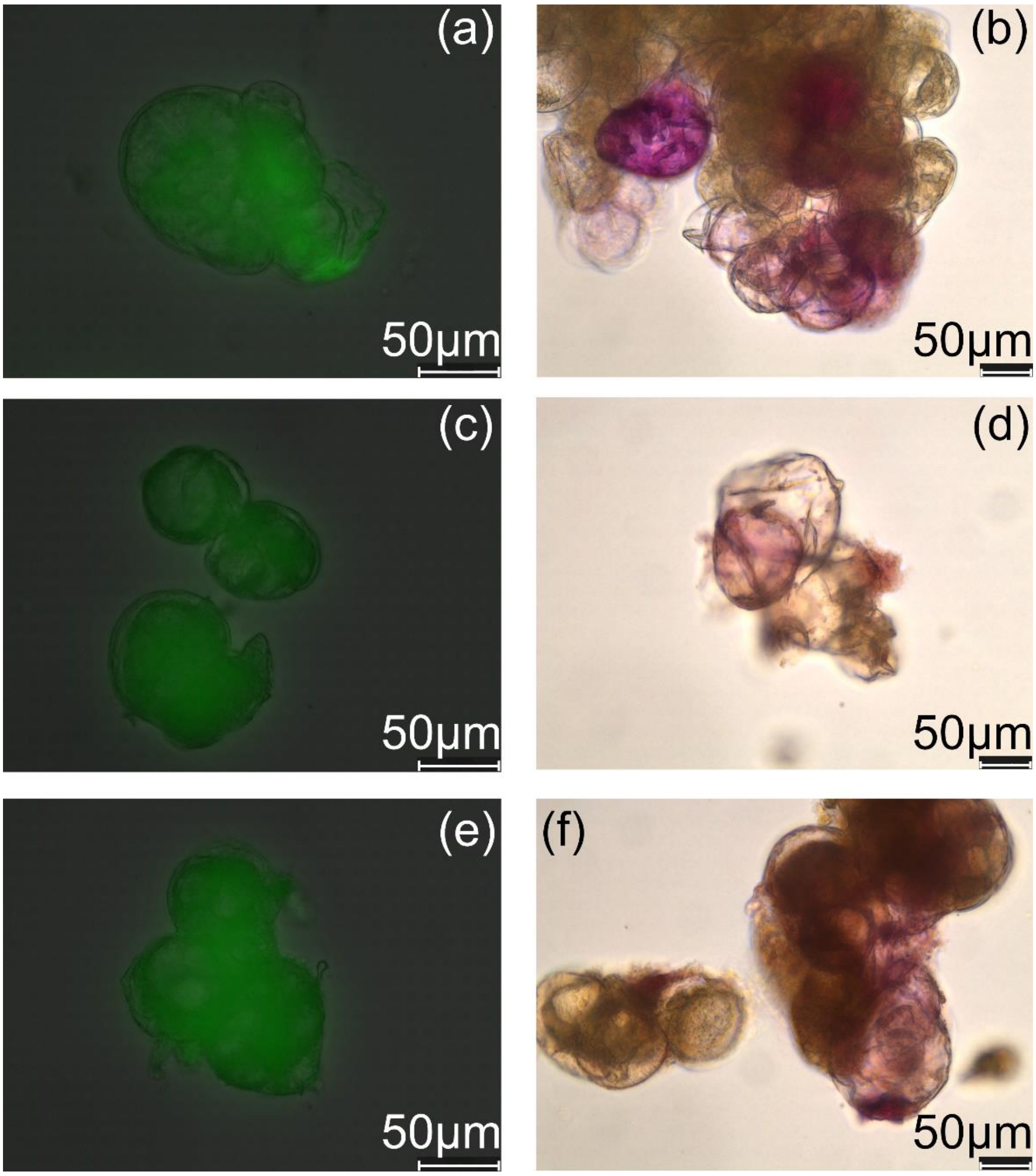


**Figure S2**: Overlay of autofluorescence and bright background (a, c, e) of cells from "ECL-inducing" liquid medium after ECL was collected, next to Wiesner test (b, d, f) of cells from the same treatments. Treatment for (a) and (b) is 0→0i, for (c) and (d) is 0→0.5i and for (e) and (f) is 0→1i. Magnification of the images is 40x and scalebars are 50 µm.

**Table S1**: Comparison of molecular weight from THF-SEC for ECL collected from "non-inducing" and "ECL-inducing" liquid media. The media marked with "i" and a star (*) are the "ECL-inducing" ones and the data are reported from previous work.[21]

| Sample | Mn (Da) | Mw (Da) | Đ |
|---|---|---|---|
| 0→1 | 2800 | 4600 | 1.63 |
| 0→1i* | 1900 ± 370 | 3200 ± 750 | 1.63 ± 0.16 |
| 1→0 | 2400 | 3800 | 1.55 |
| 1→0i* | 1700 ± 150 | 2700 ± 400 | 1.55 ± 0.11 |
| 1→1 | 2000 | 2900 | 1.47 |
| 1→1i* | 1800 ± 290 | 3000 ± 640 | 1.59 ± 0.11 |

**Table S2**: Composition of ECL collected "non-inducing" and "ECL-inducing" liquid media. The values are per 100 C9 units. The standard deviation is reported from the analysis performed on ECL collected from two separate batches, as described in the main text. It should be noted that one repetition was performed on lower concentration of ECL in deuterated solvent, which could be the reason for the different integral values. The samples marked with * are reported from our previous work.[21]

| Sample | β-O-4' | β-β' | β-5' | DBDO | Cinnamyl alcohol | H-units |
|---|---|---|---|---|---|---|
| 0→0 | 35.5 ± 0.7 | 5.5 ± 0.7 | 19.0 ± 1.4 | 3.0 ± 0.0 | 11.0 ± 0.0 | 1.4 ± 0.6 |
| 0→0i* | 31 | 10 | 18 | 5 | 8 | 2 |
| 0→1 | 32 ± 2.8 | 5.5 ± 0.7 | 19.5 ± 0.7 | 3.5 ± 0.7 | 14.0 ± 2.8 | 1.8 ± 0.6 |
| 0→1i* | 27 | 12 | 16 | 6 | 14 | 3 |
| 1→0 | 37.5 ± 4.9 | 5.0 ± 0.0 | 19.5 ± 0.7 | 3.0 ± 0.0 | 9.5 ± 3.5 | 1.6 ± 1.2 |
| 1→0i* | 26 | 10 | 14 | 2 | 16 | 5 |
| 1→1 | 31.0 ± 0.0 | 6.0 ± 1.4 | 19.5 ± 2.1 | 2.5 ± 0.7 | 11.0 ± 5.7 | 1.7 ± 0.9 |
| 1→1i* | 29 | 8 | 15 | 6 | 10 | 6 |
